# Discovery of host-directed modulators of virus infection by probing the SARS-CoV-2-host protein-protein interaction network

**DOI:** 10.1101/2022.06.03.494640

**Authors:** Vandana Ravindran, Jessica Wagoner, Paschalis Athanasiadis, Andreas B. Den Hartigh, Julia M. Sidorova, Aleksandr Ianevski, Susan L. Fink, Arnoldo Frigessi, Judith White, Stephen J. Polyak, Tero Aittokallio

## Abstract

The ongoing coronavirus disease 2019 (COVID-19) pandemic has highlighted the need to better understand virus-host interactions. We developed a network-based algorithm that expands the SARS-CoV-2-host protein interaction network and identifies host targets that modulate viral infection. To disrupt the SARS-CoV-2 interactome, we systematically probed for potent compounds that selectively target the identified host proteins with high expression in cells relevant to COVID-19. We experimentally tested seven chemical inhibitors of the identified host proteins for modulation of SARS-CoV-2 infection in human cells that express ACE2 and TMPRSS2. Inhibition of the epigenetic regulators bromodomain-containing protein 4 (BRD4) and histone deacetylase 2 (HDAC2), along with ubiquitin specific peptidase (USP10), enhanced SARS-CoV-2 infection. Such proviral effect was observed upon treatment with compounds JQ1, vorinostat, romidepsin, and spautin-1, when measured by cytopathic effect and validated by viral RNA assays, suggesting that HDAC2, BRD4 and USP10 host proteins have antiviral functions. Mycophenolic acid and merimepodib, two inhibitors of inosine monophosphate dehydrogenase (IMPDH 1 and IMPDH 2), showed modest antiviral effects with no toxicity in mock-infected control cells. The network-based approach enables systematic identification of host-targets that selectively modulate the SARS-CoV-2 interactome, as well as reveal novel chemical tools to probe virus-host interactions that regulate virus infection.

**Synopsis:** 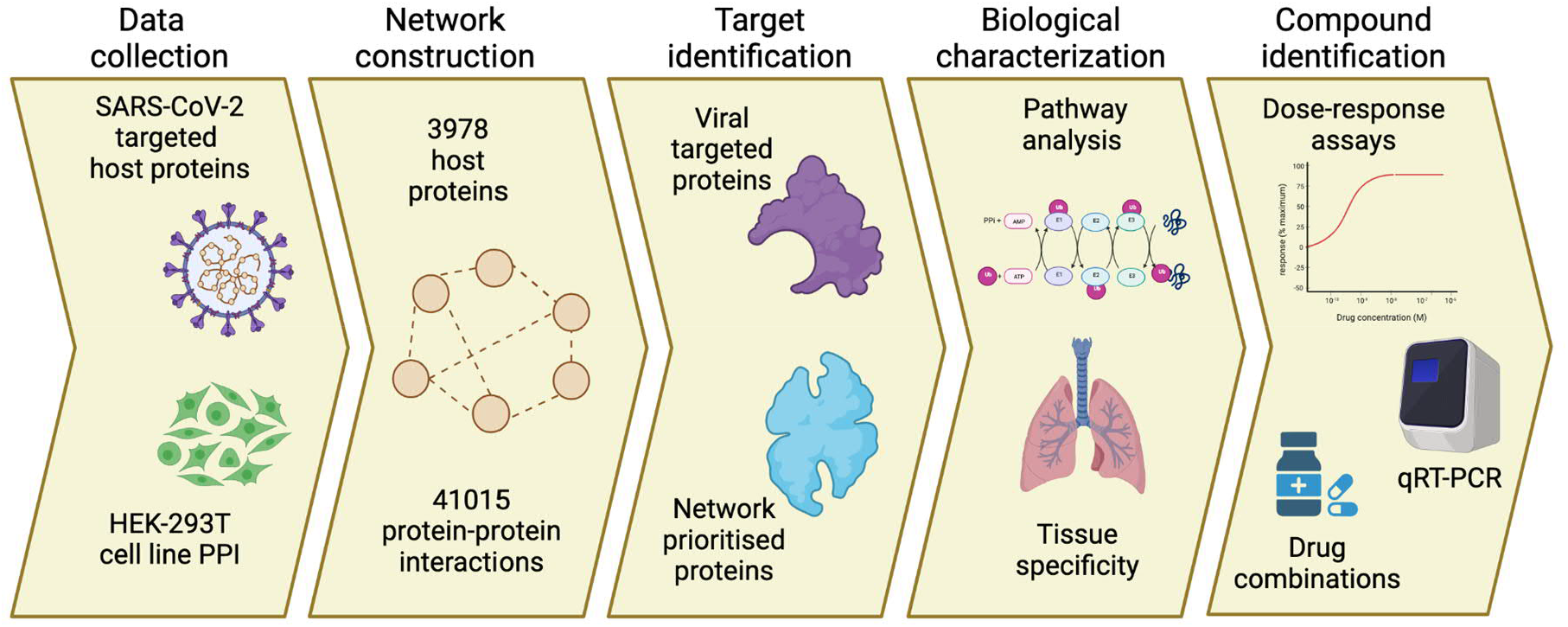

Viruses exploit host machinery and therefore it is important to understand the virus-host dependencies to gain better insight of the key regulators of viral infection.

- Using a context-specific SARS-COV-2 PPI network, a computational framework was developed to identify host modulators of viral infection.
- Chromatin modifying host proteins HDAC2 and BRD4, along with deubiquitinating enzyme USP10, act as antiviral proteins.
- IMPDH inhibitors mycophenolic acid and merimipodib showed modest antiviral response to SARS-COV-2 infection, and no toxic effects.
- Cell context specificity is a critical factor when identifying selective modulators of viral infection and potential antiviral therapeutics.
- Topology-based network models cannot distinguish between host-proteins, the inhibition of which leads to either virus suppressive or enhancing effects.

## INTRODUCTION

The global outbreak of severe acute respiratory syndrome coronavirus-2 (SARS-CoV-2) requires the development or re-positioning of effective and safe therapies against the virus and coronavirus disease 2019 (COVID-19). In particular, drug repurposing enables a fast track to identify small-molecules that target emergent viruses, including SARS-CoV-2, by focusing on viral targeted host proteins and their cellular pathways as targets for the therapeutic intervention. In response to infection, host cells mount antiviral responses. However, SARS-CoV-2, like many other viruses, has developed various strategies to escape cellular antiviral responses, similarly to Dengue, SARS-CoV-1, MERS-CoV (1). It is therefore important to elucidate the interactions of the SARS-CoV-2 with host proteins to gain better insights into virus infection and pathogenesis, and to potentially reveal novel options for therapeutic intervention.

Recent large-scale proteomic and genomic profiling studies have elucidated various mechanisms by which SARS-CoV-2 alters host cells (2-5). Several studies have utilised virus-host protein-protein interactions (PPIs) to construct networks of SARS-CoV-2–induced pathways and to uncover novel targets and potential host-targeting repurposing agents (6-8). Some of these network-based studies have also provided a comprehensive resource of the biological pathways and potential drugs for the SARS-CoV-2 targeted host proteins (9-11). However, many of the small molecules identified through computational network studies have not been experimentally validated, and it has been argued that a truly impactful computational method should lead to actionable and experimentally testable hypotheses to enable the discovery of novel compounds or commbinations (12). For instance, Gysi et al. experimentally screened 918 compounds in VeroE6 cells and found that none of the predictive algorithms offers consistently reliable outcomes across all metrics and against investigational compounds in clinical trials (13). Another challenge in experimental testing of model predictions is the high cell context-specificity of the compound responses, often making the screening results based on a single cell line challenging to reproduce in other relevant cell types.

Here, we developed a computational-experimental approach that integrates virus-host protein interactions with both a human PPI network as well as tissue expression profiles to (i) prioritise host proteins that can be targeted by compounds in clinical development and (ii) to experimentally probe how these compounds modulate virus infection. The key feature of our integrated approach is that it exploits the cell-context specific dependencies of the virus on specific host proteins and pathways during replication. To expand the host targets, biological pathways, and compound spaces, we used a random walk with restart, a probabilistic network propagation algorithm, to identify additional host targets from the extended set of nearest neighbours of virus-interacting proteins (VIPs). We explored the biological function and targeting effect of such network-inferred proteins (NIPs), which may not interact directly with viral proteins, but can be functionally related to VIPs through various pathways within the SARS-CoV-2 interactome.

We tested selective inhibitors of the identified host proteins in two cell-lines (HEK293T cells expressing ACE2 and TMPRSS2 and lung epithelial Calu 3 cells) with multi dose assays to understand the dose-and context-specific effects of the inhibitors on viral infection in relevant cell models. Interestingly, we identified several host-targeting compounds that enhance virus infection, suggesting that the target proteins act as antivirals. Furthermore, two host-targeting compounds showed modest suppressive effects on virus infection, and no toxic effects in mock-infected control cells, demonstrating that even though the network-based approach identified therapeutically safe targets, it cannot distinguish between the virus suppressive and enhancing effects. Collectively, our computational-experimental approach supports the discovery of novel host-targeting modulators of virus infection, as well as novel chemical tools for probing how virus-host interactions regulate virus infection.

## RESULTS

### Viral interacting proteins are critically positioned for network information flow

Similar to other viral infections, the SARS-CoV-2 life cycle is mediated through a complex system of protein interactions, modelled here as network graphs, with cellular proteins depicted as nodes and their interactions as edges. To model the system-wide virus-host interactions, we used a PPI network among 298 human proteins identified previously as having interactions with SARS-CoV-2 virus proteins. Since the initial characterization of virus interacting host proteins (VIPs) was carried out in the HEK293T cell line (14), we constructed our SARS-CoV-2 PPI network in this same cellular context by mapping the interactions between the VIPs and their nearest neighbours derived from pull down experiments reported in the Bioplex interactome (15). The HEK293T PPI network consists of 3,978 protein nodes and 41,015 interaction edges, and it has two connected components: a giant component of 3,975 nodes and a smaller component of 3 nodes (**Supplementary Table S1a**). In the downstream network analyses, we focused on the giant component that contains 297 VIPs and their 3678 nearest neighbour proteins (non-VIPs) that are not targeted directly by the viral proteins (**Figure 1A**).

**Figure 1.**
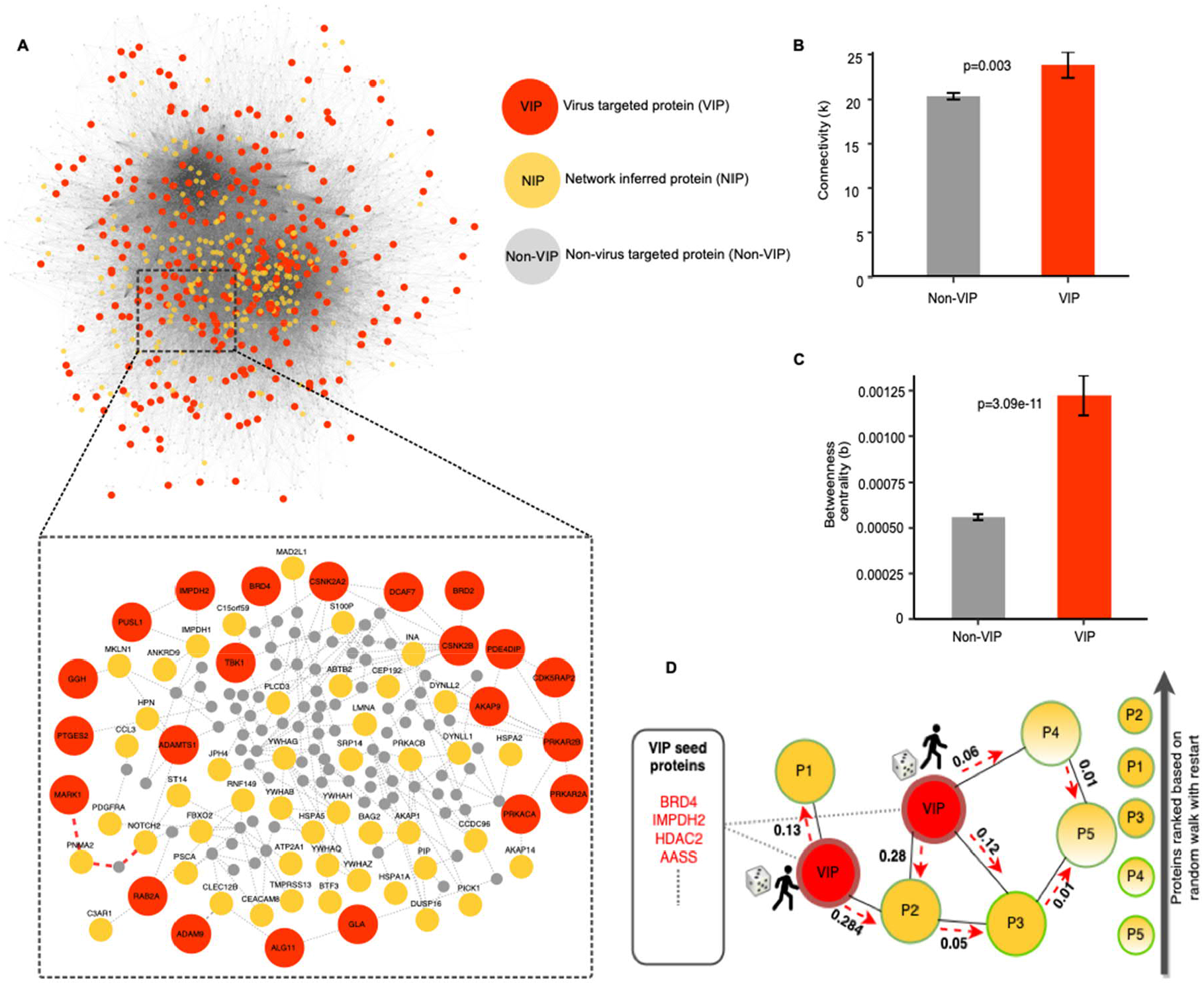
Topological network analysis of the SARS-CoV-2 PPI network. (A) The giant component of the SARS-CoV-2-host PPI network that contains 297 VIPs and 3678 non-VIPs. The inset shows a subnetwork of select VIP nodes. The connections marked in red show the shortest path distance from the MARK1 VIP node to NOTCH2 node, as an example of calculating the neighbourhood size from VIP to non-VIP nodes (neighbourhood size of 3). (B) Degree, i.e., the number of connections, for VIP and non-VIP nodes (the bars show mean ± SEM). (C) Betweenness centrality, i.e., the number of shortest paths through a given node, for VIP and non-VIP nodes (mean ± SEM). (D) Operation of the random walk with restart (RWR) algorithm. The red seed nodes are the set of 298 VIPs from which the random walker starts exploring the network (marked by red arrows). After iterating through all the nodes in the network, a probability score is assigned to each node in the network, ranked from the highest to the lowest probability, which was used to identify the 200 top-ranked NIP nodes.

To investigate the role of the host proteins targeted by the virus in the PPI network, we calculated several network information measures that quantify the topological inter-connectivity of the giant network component (**Supplementary Table S1b**). The connectivity (*k*), i.e., the number of direct interacting partners of each protein, showed that the SARS-CoV-2 virus host VIPs have a higher connectivity (mean 23.82), compared to the non-virus targeted host proteins (non-VIPs) (mean 20.36, P=0.0026, Wilcoxon test; **Figure 1B**). The central position of VIPs in the network information flow became even more pronounced in terms of the betweenness centrality (*b*), i.e., the number of shortest paths between each pair of nodes that pass-through a given node, which showed that VIPs have significantly higher centrality (mean 0.0012), compared to non-VIPs (mean 0.00055, P=3.084×10^−11^, Wilcoxon test; **Figure 1C**). This observation is consistent with other studies that have shown that viruses tend to target network hubs and bottlenecks in the PPI network (16,17), since viruses and host proteins are constantly competing for binding partners that interact with proteins in various pathways during the infection phase.

### Identification of additional host proteins that are indirectly regulated by VIPs

To further study the information flow through VIPs, we characterised their neighbourhood sizes, i.e., the length of the shortest paths from a VIP to any non-VIP nodes in the giant component of the network. We found that most VIP nodes can be reached by traversing as few as 3 or 4 edges from any other non-VIP node. This is comparable to the average path length from a random node to any other node in the network (mean 3.422, **Figure S1**). This topological network analysis provides information for the positioning of host-directed modulators of virus infection that are either directly interacting with viral proteins (VIPs), or are components of viral-regulated pathways, where proteins do not directly interact with viral proteins (non-VIPs), but are nonetheless required for viral replication. In particular, the central position of VIPs in the network makes it critical to investigate the potential side effects of inhibiting these host targets, as network hubs often interact with many other proteins involved in normal cellular processes and biological pathways, targeting of which may lead to toxic effects on non-infected cell (18).

The above network analysis suggests that it is also important to consider host proteins that are not directly targeted by viral proteins yet are part of pathways regulated by viruses. For such extended host target mapping, we employed random walk with restart (RWR), a probabilistic network propagation algorithm (19), which was applied here in the identification of proteins connected to VIPs (see Materials and Methods). The RWR algorithm explores the network vicinity and function of the protein seed set (here, VIPs), based on the premise that proteins with similar functions tend to lie close to each other in the PPI network. Thus, the RWR algorithm identifies proteins within the network that may be functionally similar to the VIPs by topologically scanning each node in the PPI network from the VIP seed set, and then assigns a probability score to the non-seed proteins (**Figure 1D**). The proteins were ranked based on the probability assigned by the RWR algorithm, so that the higher the probability, the closer the proteins are to the VIPs. In this way, we shortlisted 200 top-ranked proteins, and termed them as network-inferred proteins (NIPs) (**Supplementary Table S1b**) **(Figure 1A)**.

### Biological pathway characterization of the identified host targets

To investigate whether the network-inferred targets (NIPs) also share similar biological function with the VIPs, as suggested by their network similarity, we performed gene set enrichment analysis (GSEA), separately for the VIP and NIP targets, and then compared the biological pathways and gene ontology (GO) terms between the two target sets. A total of 11 biological processes were similarly enriched in the NIP and VIP sets, including protein folding (P=6.70×10^−8^, 1.43×10^−7^), establishment of protein localisation to membranes (P=6.80×10^−8^, 2.90×10^−6^), and protein targeting (P=6.59×10^−6^, 2.62×10^−8^) (**Figure 2**). Apart from these biological processes that are known to be involved in viral processing (20,21), we also identified several additional common biological processes between the two host target sets that related to immune responses, such as neutrophil activation, mediated immune responses and degranulation (**Supplementary Table S2**). We further observed that the network-predicted NIP targets also shared other relevant pathways with the VIP set in terms of KEGG pathways. For instance, the NIP and VIP targets were both enriched for proteins involved in processing in the endoplasmic reticulum (ER) pathway (P=1.198×10^^-5^, 3.72×10^^-5^; **Figure 2**), which is biologically plausible as the ER is involved in viral replication and assembly (22).

**Figure 2.**
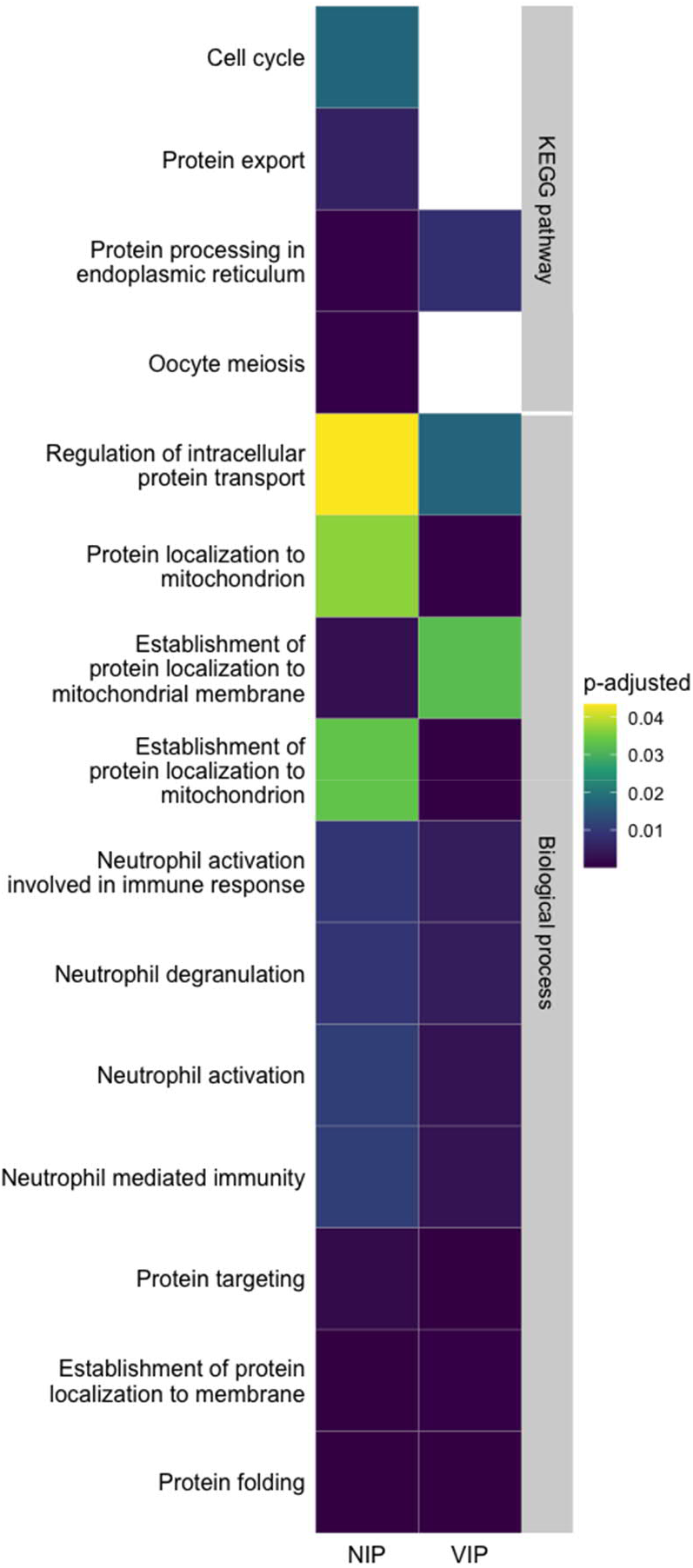
GO biological process and KEGG pathways that are commonly or uniquely enriched among the NIP and VIP targets. The enrichment P-values were corrected for multiple testing with Benjamini-Hochberg test, and the pathways with adjusted P<0.05 were considered significant.

Interestingly, the NIP target set also extended biological processes beyond those captured by the VIP targets. For instance, NIP targets were enriched for the cell cycle pathway (P=3.62×10^−4^), which highlights its importance in viral replication (23,24). These network-based results also support previous studies that have shown how SARS-CoV-2 hijacks the rough endoplasmic reticulum (RER)-linked host translational machinery for its replication (11,30). In addition, NIP proteins were enriched for pathways related to protein export (P=8.99×10^−5^), which also includes secretory pathways that are hijacked by the virus to carry out their essential functions, such as virus replication, assembly, and egress, demonstrating that viruses evade host cellular pathways to promote their propagation (25). These results indicate that the extended target set consisting of the top-200 NIP host proteins identified by the RWR algorithm not only populate similar pathways as VIPs but are also implicated in various biological processes related to viral infection.

### In silico identification of host-targeted compounds that modulate viral infection

To disrupt the SARS-CoV-2 interactome, we sought potent and selective compounds that inhibit the identified host proteins (both VIPs and NIPs). We prioritised drugs that are approved for other indications and investigational compounds currently being tested in clinical trials (phase 1-3), instead of preclinical candidates that would take longer time to develop as COVID-19 modulators. We first identified compounds that inhibit the identified host targets (VIPs and NIPs) through in-silico screening using target activity data from the ChEMBL database (26). For the VIP targets, 6,458 unique compounds were identified that cover 27 of the 298 targets (9%; **Figure 3A**). For the NIP targets, 2,754 unique compounds were identified that cover 25 of the top-200 targets (13%; **Figure 3B**). We considered compounds with a bioactivity less than 1000nM against the host targets as potent inhibitors, and if multiple compounds were found to inhibit a particular target, we selected the inhibitor with the highest potency for the experimental validations.

**Figure 3.**
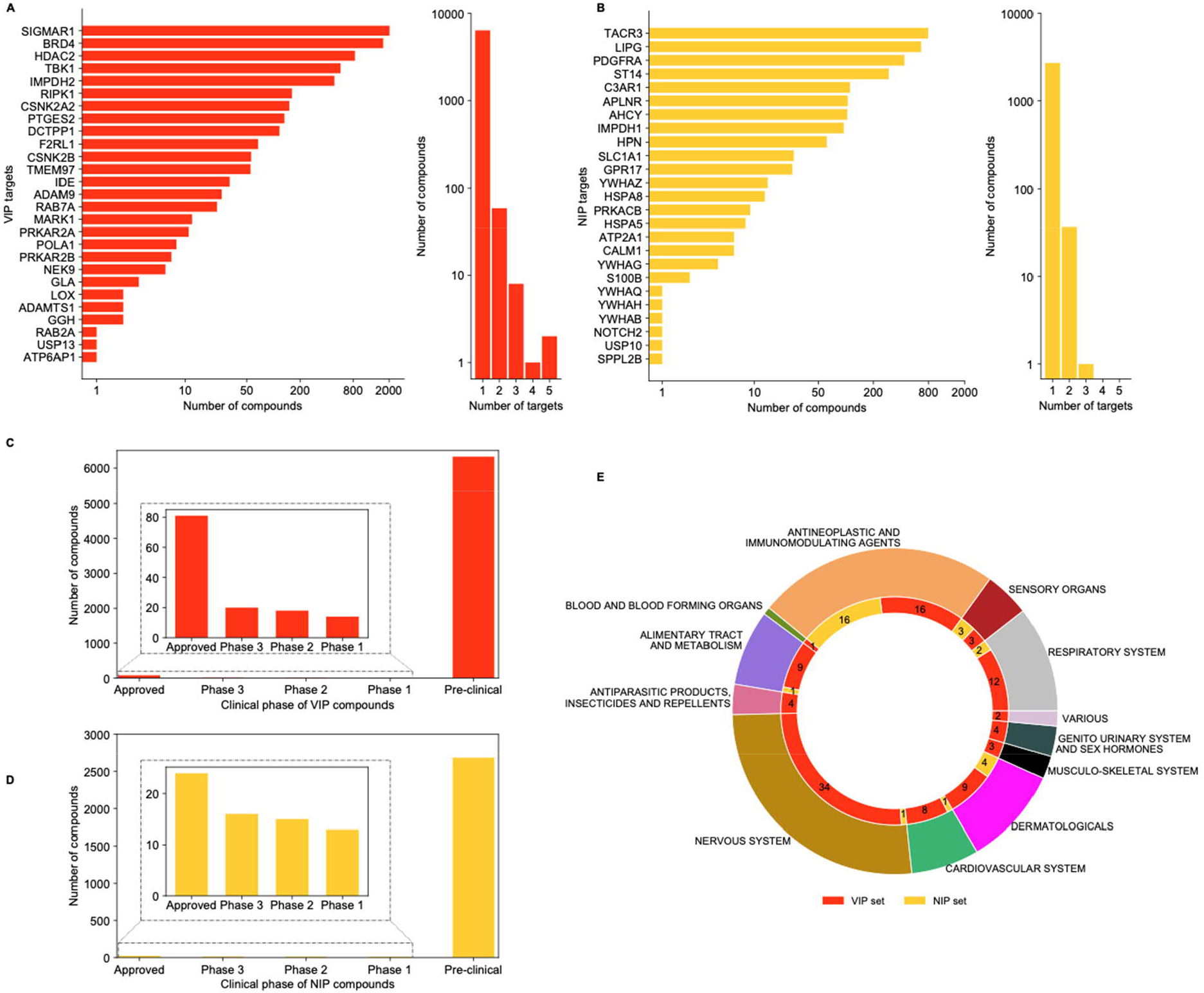
Identification of potent compounds that inhibit the host target proteins based on ChEMBL database (bioactivity less than 1000nM, see Materials and Methods). (A) The number of compounds identified for the 27 VIP targets (left histogram), and the number of VIP targets for each potent compound (right histogram). (B) The number of compounds identified for the 25 NIP targets (left), and the number of NIP targets for each potent compound (right). (C) Clinical development phase of compounds that target VIPs. (D) Clinical development phase of compounds that target NIPs. (E) Anatomical Therapeutic Chemical (ATC) classification of the 101 approved compounds that target select VIPs and NIPs.

Considering the number of compounds for VIP and NIP targets, many of the identified proteins are targeted by multiple compounds, suggesting that most of these host targets are already well-studied in drug discovery (**Figure 3A,B**, left distributions). The distribution of compounds per target between the VIPs and the NIPs were relatively similar (P=0.75, Kolmogornov-Smirnov test). Notably, most of the compounds have been reported to show activity against a single target only among these target sets, suggesting that the inhibitors may be selective against the host targets in each set (**Figure 3A,B**, right distributions). However, most of the compounds that selectively target VIPs and NIPs are still in a pre-clinical phase (**Figure 3C,D**). This systematic compound screen resulted in a total of 9,079 potent inhibitors of the 52 host proteins (27 VIPs and 25 NIPs) (**Supplementary Table S3**), out of which only 101 (0.01%) are currently approved for other indications (**Figure 3E**), i.e., representing potential repurposing opportunities.

### Selection of a subset of host targets that is most relevant for COVID-19

To select the host targets and their inhibitory compounds for experimental validation, we further shortlisted our target lists from both the VIP and NIP sets, based on the expression levels of the host proteins in cells relevant for COVID-19 using expression data from the Human Protein Atlas (27,28). Specifically, we chose to use the target expression in lung epithelial cells and cells of the upper and lower respiratory tract, since the virus infects the upper respiratory tract, causing flu-like symptoms, and the lower respiratory tract, causing severe respiratory disorders.

Among the VIP targets, we found that BRD4 (bromodomain and extraterminal (BET) protein 4), RAB7A (Ras-related protein 7A) and HDAC2 (Histone deacetylase 2) were relatively highly expressed in respiratory epithelial cells of the nasopharynx and bronchus, as well as in lung macrophages and alveolar cells **(Figure 4)**. Similarly, IMPDH2 (Inosine-5-monophosphate dehydrogenase 2) showed high expression in lung macrophages and medium expression in respiratory epithelial cells of the nasopharynx and bronchus. BRD4 is a member of the Bromo and Extra Terminal (BET) family of proteins that bind acetylated histones and play roles in regulation of transcription, DNA repair, and replication (29,30). In transcription, BRD4 is involved in activation of multiple genes, including those central to proinflammatory and host immune responses (31-33).

**Figure 4.**
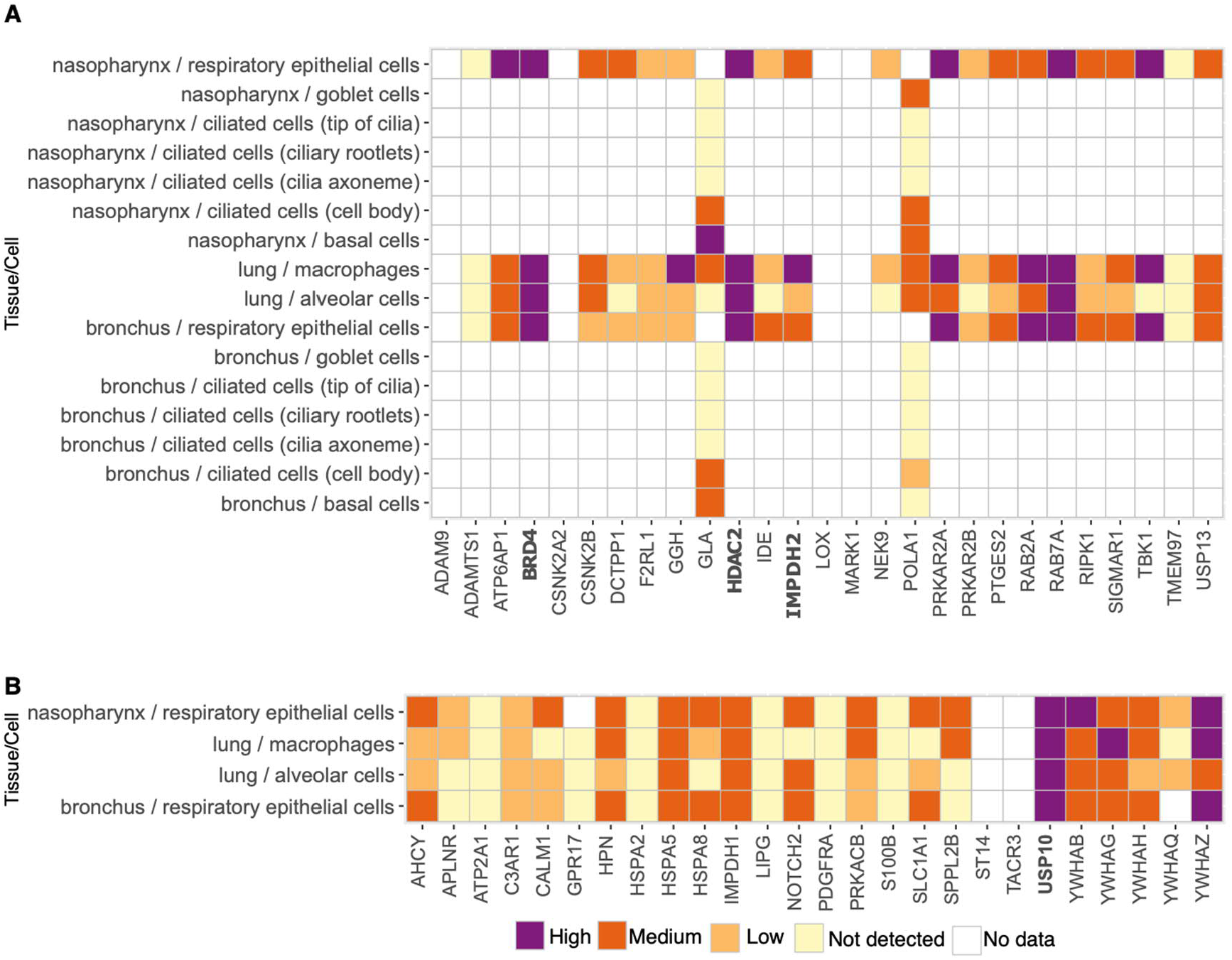
Protein expression in cells of upper and lower respiratory tract across the (A) 27 VIP targets and (B)25 NIP targets. The expression classes are from the Human Protein Atlas (colour legend).

HDAC2, a class I histone deacetylase, regulates transcription and other DNA metabolic processes by removing the epigenetic marks of open chromatin – acetyl post-translational modifications – from histones H3 and H4 (34). Along with other HDAC family members, HDAC2 is an important player in numerous cellular processes (35), and its downregulation is associated with hyperinflammatory conditions (36). HDAC2 can act as either a restriction or dependency factor for various viruses. For instance, HDAC2 acts as a dependency factor for herpes simplex viruses (HSV) (14,37), and inhibition of HDAC2 reduces human adenovirus expression and replication (38). On the other hand, HDAC2 contributes to the antiviral response against influenza viruses (39).

IMPDH2 regulates NF-κB signalling, and SARS-CoV-2 infection leads to NF-κB activation and induction of proinflammatory cytokines (40). IMPDH2 also plays a role in purine biosynthesis, which is needed for viral RNA replication (41). While RAB7A (Ras-related protein 7A) had high expression in lung macrophages and in respiratory epithelial cells of the bronchus **(Figure 4A)**, it was de-prioritised because of its key role in the maturation of late endosomes, which are not the primary sites through which SARS-CoV-2 enters human lung cells; most SARS-CoV-2s enter lung cells through fusion at the plasma membrane (42), while for the Omicron variants of concern (VOC), the virus appears to also use the endosomal route (43,44).

Among the NIP targets, USP10 (Ubiquitin specific peptidase 10) showed high expression in respiratory epithelial cells of the nasopharynx and bronchus, and in macrophages and alveolar cells of lung tissue (**Figure 4B**). Similarly, YWHAZ (Tyrosine 3-Monooxygenase/Tryptophan 5-Monooxygenase Activation Protein Zeta) was highly expressed in respiratory epithelial cells of the nasopharynx and bronchus and in lung macrophages, and it had a medium expression in lung alveolar cells. Even though only preclinical inhibitors of these NIP host protein targets are currently available, we nonetheless pursued USP10 in the experimental validation, as this protein plays a role in stress granules and RNA processing, storage, and degradation, that are used by flaviviruses such as West Nile and Dengue virus (45).

Based on the expression analyses of the protein targets in the VIP and NIP sets, we shortlisted BRD4 that is targeted by the SARS-CoV-2 envelope protein E, along with HDAC2 and IMPDH2 that are targeted by the viral protease (NSP-5) and exoribonuclease (NSP-14), respectively (**Table 1**). Among NIPs, we focused on USP10, which engages the host VIP proteins G3BP1 and G3BP2 that directly engage the viral N protein in the experimental validation.

**Table 1.** List of VIPs and NIPs shortlisted for the experimental assessment of compound effects.

| Protein class | Host protein | Viral protein | Degree | Betweenness centrality | Implicated in other viruses | Associated role |
| --- | --- | --- | --- | --- | --- | --- |
| Epigenetic regulators | BRD4 | E | 26 | $1.46 \times 10^{-3}$ | HPV -1/2/5/7/9/10, EEPV4, HHV8, CRPV2 | Regulate genes crucial for cell cycle, progression, inflammation and immune response |
| | HDAC2 | Nsp5 | 27 | $7.99 \times 10^{-4}$ | Adeno-associated dependoparvovirus A, HPV, FLUAV | Plays an important role in transcriptional regulation, cell cycle progression and developmental events |
| Enzymes | IMPDH2 | Nsp14 | 7 | $1.07 \times 10^{-4}$ | Dengue virus, HSV-1, HIV-1, FLUAV, Measles morbillivirus | Plays an important role in the regulation of cell growth and purine metabolic processes |
| | USP10* | N | 8 | $1.75 \times 10^{-4}$ | HHV 8, FLUAV, Zika virus | Plays a role in regulation of autophagy |

\*This NIP interacts with VIP G3BP1/G3BP2

### Experimental assessment of the host target inhibition on virus infection

Our computational prioritisation approach identifies host targets that are expected to modulate virus infection within the SARS-CoV-2-host PPI network; however, similar to other network-based models, it does not predict whether the inhibition of the host targets leads to either virus suppressive or enhancing effects, nor whether the host target inhibition results in any toxic side effects. Therefore, in the experimental validations, we tested not only the effects of the host-targeting compounds on SARS-CoV-2 infection, but also their effects on the viability of mock-infected control cells. We experimentally tested seven selected compounds that have inhibit the VIPs BRD4, IMPDH2, and HDAC2, along with spautin-1, which inhibits the NIP USP10 and the VIP USP13 (**Table 2**). The compounds were first tested for modulation of SARS-CoV-2 infection in HEK293T cells that overexpress the key SARS-CoV-2 host factors ACE2 and TMPRSS2 (46). Confirmatory experiments were performed in Calu 3 human lung cells (47), which are well-established targets for SARS-CoV-2 (42).

**Table 2.**
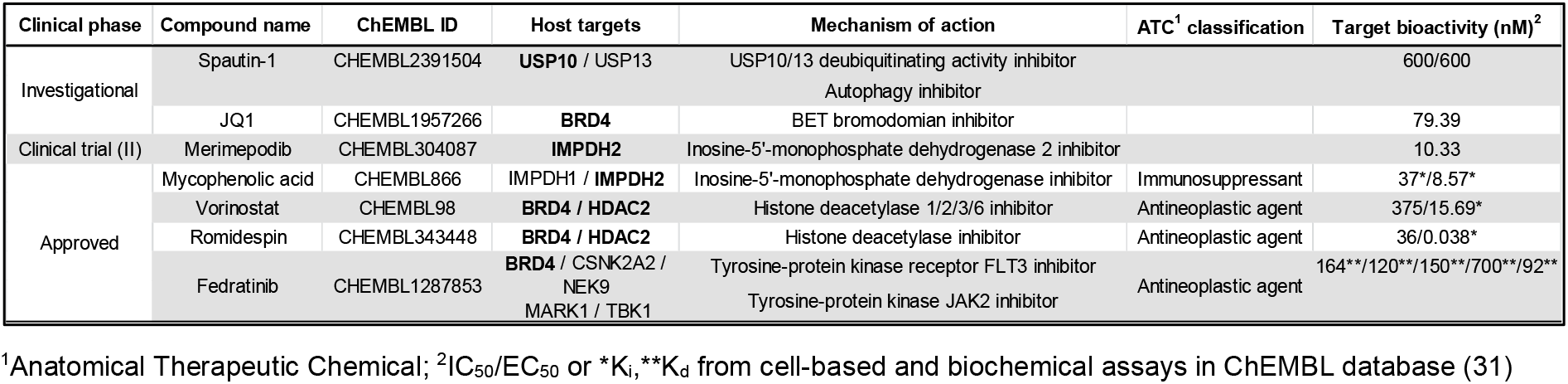
Compounds tested in 293T-ACE2-TMPRSS2 and Calu 3 cells in the present work. The boldfaced host targets were identified by the network-approach.

### Discovery of host modulators that enhance SARS-CoV-2 infection

We noticed that several compounds that target chromatin regulating proteins, such as BRD4 and HDAC2 and deubiquitinating enzymes USP10 and USP13, did not show any significant antiviral activity in our disease system (**Figure 5**, efficacy curve). In the virus-infected cells, inhibition of these proteins actually increased virus-induced cytopathic effect (CPE), when compared to the DMSO control. This increased CPE was reflected in a negative antiviral efficacy (see Materials and Methods), and suggest that these compounds enhanced virus infection. This enhanced CPE (pro-viral effect) occurred at concentrations that were not directly cytopathic to cells (**Figure 5**, viability curve), indicating moderate toxic effects of the compounds in the non-infected control cells that were included in each assay to assess effects of compound on cell viability.

**Figure 5.**
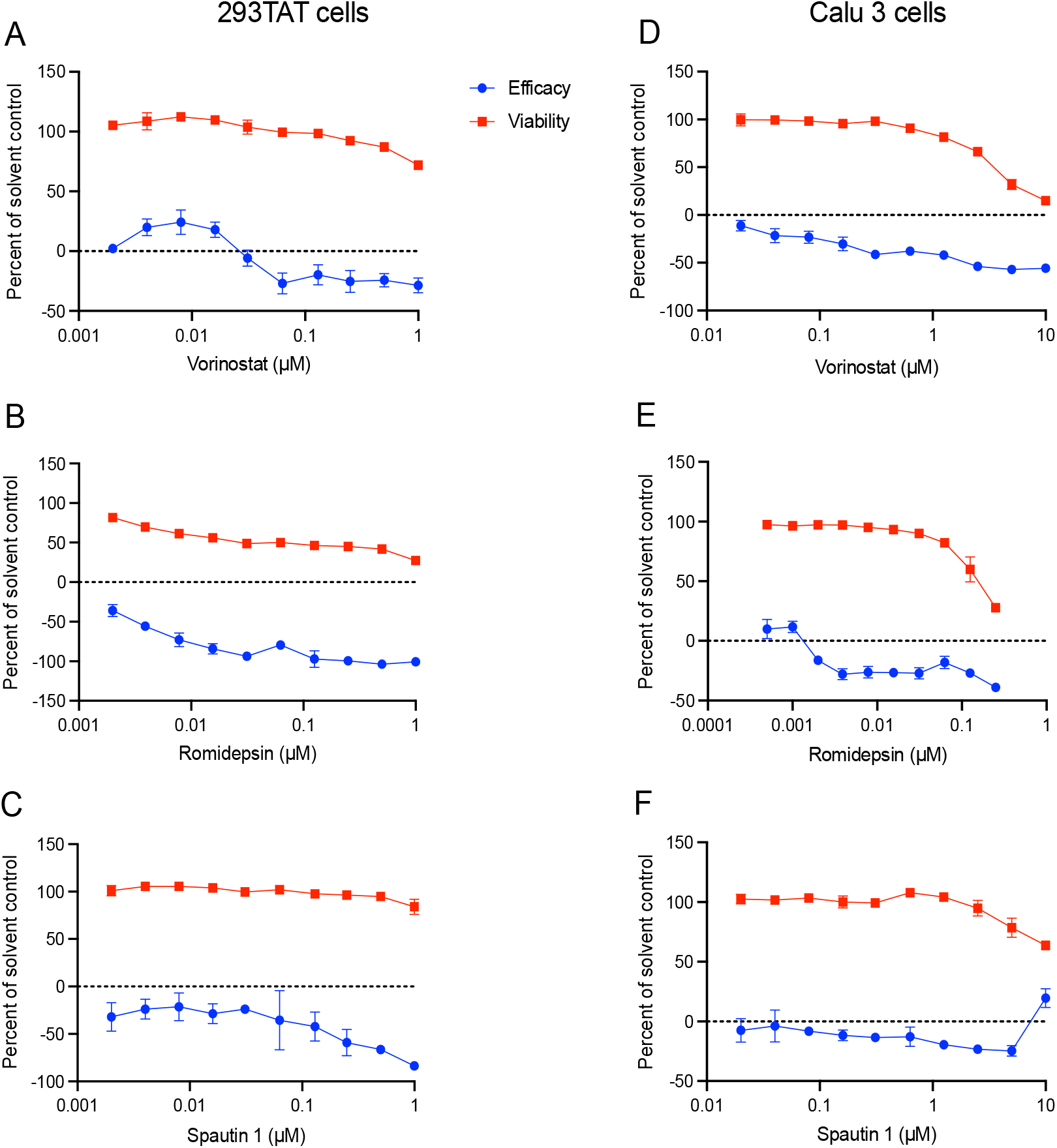
Experimental assessment of the host target inhibition. (A-C) Compounds that inhibit HDACs and USP10/13 enhance CPE during virus infection in 293TAT cells (i.e., appear to have pro-viral effects). Cells were treated with the indicated concentrations of compounds immediately prior to infection with SARS-CoV-2 at a MOI of 0.01. Parallel wells contained cells treated only with compounds to study toxicity. Forty-eight hours post infection, cell viability was assessed using the CellTiter-Glo assay and effects on virus infected cells (efficacy) and non-infected cells (viability) were calculated as described in Materials and Methods. Data points reflect average and standard deviations of triplicate experiments per condition. (D-F) Confirmatory assays in Calu 3 cells. Cells were treated with the indicated concentrations of compounds for 2 hours prior to infection with SARS-CoV-2 at a MOI of 0.1. Parallel wells contained cells treated only with compounds. Ninety-six hours post infection, cell viability was assessed using the CellTiter-Glo and effects on virus infection (efficacy) and viability were calculated as described in Materials and Methods. Data points reflect average and standard deviations of triplicate experiments per condition.

Compounds that showed such pro-viral phenotype included vorinostat, a broad range inhibitor of histone deacetylases (HDACs) (**Figure 5A**), and romidepsin, a more selective HDAC inhibitor that targets HDAC1 and HDAC2 (**Figure 5B**). Similarly, the investigational compound, spautin-1, which inhibits the NIP USP10 and VIP USP13, enhanced CPE at concentrations that were not inherently toxic to non-infected cells (**Figure 5C**). To further investigate this putative proviral effect, we tested these compounds for CPE during SARS-CoV-2 infection also in Calu 3 human lung cells, where the compounds again enhanced CPE during virus infection at concentrations that were not toxic to non-infected Calu 3 cells (**Figure 5 D-F**).

If these compounds enhance virus infection, one would expect they similarly enhance viral RNA expression in the infected cells. We therefore measured the level of SARS-CoV-2 RNA expression by qRT-PCR at 24 hours post-infection of 293TAT cells, and parallel plates were assayed for CPE at 48 hours as per our protocol. The qRT-PCR assays showed that both spautin-1 and vorinostat conferred an apparent pro-viral effect as they increased the abundance of viral RNA (**Figure 6A**), consistent with their enhancement of infection seen in the CPE assay (**Figure 6B**). These data confirm that spautin-1 and vorinostat enhance virus infection, and have pro-viral effects, and suggest that their host targets may therefore have antiviral function.

**Figure 6.**
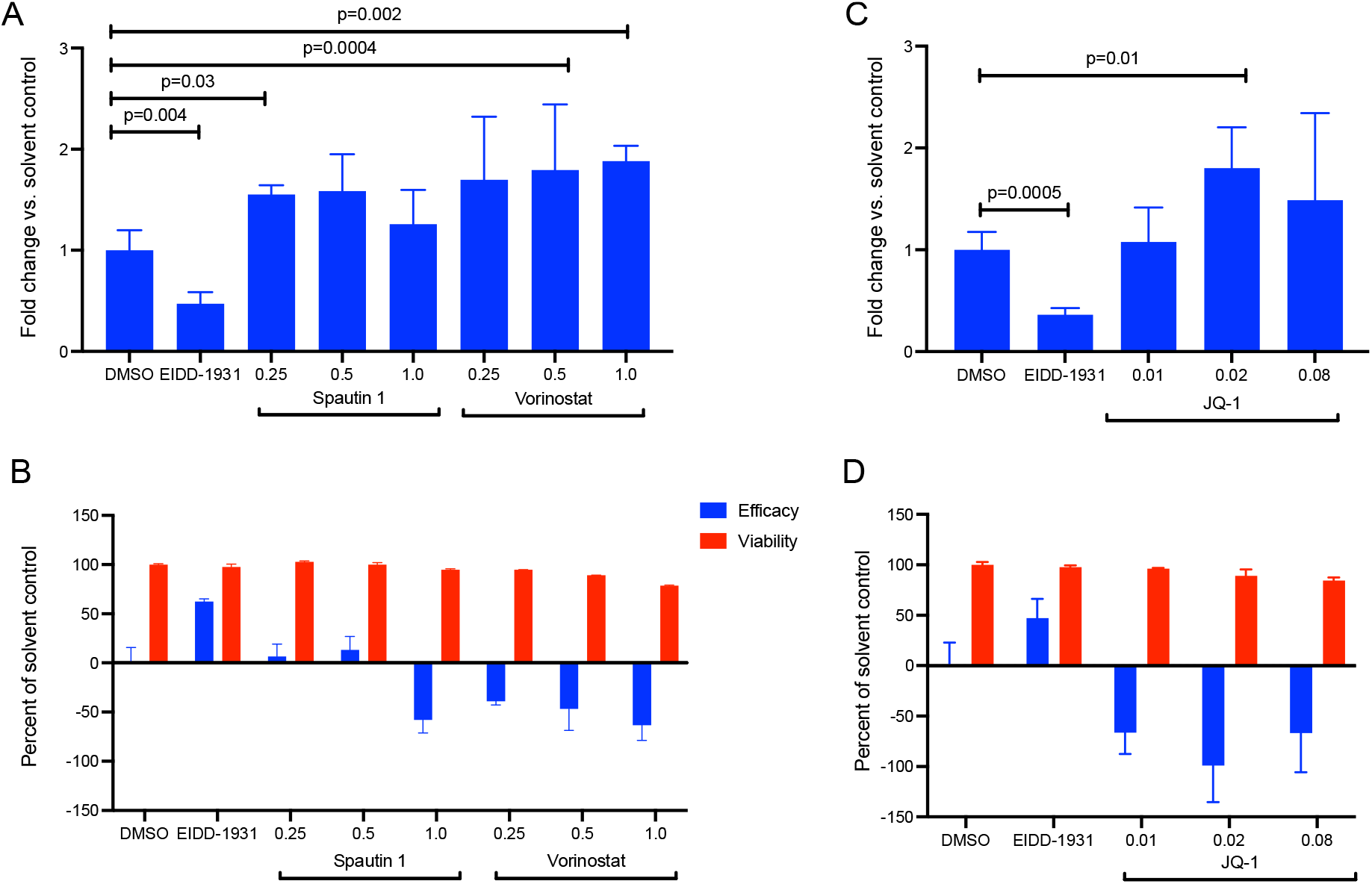
Spautin 1, vorinostat and JQ1 enhance SARS-CoV-2 infection. 293TAT cells were treated with the indicated concentrations of compounds immediately prior to infection with SARS-CoV-2 at a MOI of 0.01. Parallel wells contained cells treated only with compounds. Twenty-four hours post-infection, RNA was extracted from one plate and the relative abundance of viral RNA was determined by qRT-PCR (panels A and C). Forty-eight hours post infection, cell viability was assessed using the CellTiter-Glo assay and effect on virus infected cells (efficacy) and non-infected cells (viability) were calculated (panels B and D). Data points in panels A and B reflect average and standard deviations of triplicate and quadruplicate experiments per condition, respectively. Statistical testing was done with two-tailed Student’s t-test.

To further study the pro-viral effects of epigenetic regulators, we next examined the BRD4 inhibitor JQ1 in our CPE and viral RNA assays during infection of 293TAT cells. Similar to the other chromatin modifying enzyme inhibitors, JQ1 increased SARS-CoV-2 RNA expression and conferred a proviral effect (**Figure 6C-D**). Collectively, these observations suggest that alterations in chromatin regulation may regulate susceptibility for SARS-CoV-2. Interestingly, fedratinib, a selective oral JAK2 inhibitor recently approved in the United States for treatment of adult patients with myelofibrosis (48), which has off-target activity against the VIPs BRD4, TBK1, MARK1 and CSNK2A2, also displayed a similar proviral effect in our disease system (**Figure S2**), consistent with the fact that fedratinib acts primarily by suppressing inflammation (49), which occurs after the initial infection. However, this proviral effect could not be confirmed by viral RNA assays in the 293TAT cells.

### Network-approach identifies context-specific inhibitors of virus infection

Since a compound’s response is often highly dependent on the cell context, it is important to consider its effects also in various cell lines and assays when determining the influence of a host target modulation on viral infection. Therefore, we cross-compared the host-targeting compounds identified by our network-based analysis with the profiling studies carried out by the National Centre for Advancing Translational Sciences (NCATS) (49) with live virus infectivity CPE assay. The NCATS has tested a collection of 9,187 compounds consisting of approved, anti-infective compounds that have been reported in literature and with target information. To date, 7.94% (730/9,187) of the compounds have been classified as antivirals based on the NCATS criteria in vero E6 cell lines (compounds with curve rank>0). A total of 260 compounds from our network-based analyses were included in the NCATS compound list. Based on the NCATS CPE assay, 18.08% (47/260) of the identified compounds were classified as antivirals (**Supplementary Table S4**). Therefore, the network approach led to a 2.3-fold improvement in success rate, compared to the 7.94% success rate of the NCATS compound list. We further cross-referenced our compounds with the antivirals reported through various publications catalogued in the Coronavirus resistance database CoV-RDB database (**Supplementary Table S4**) (50).

For instance, imatinib, an approved inhibitor of Bcr-Abl kinase inhibitor, that has off-target activity against the NIP PDGFRA, showed antiviral efficacy in VeroE6 cells and human airway epithelial (HAE) cultures (**Table 3**) (51, 52). Imatinib was advanced to clinical trials, albeit without evidence for clinical efficacy (ClinicalTrials.gov, trial numbers NCT04794088, NCT04394416) (53). Another example of effective antiviral that targets TMPRSS2 and the NIPs HPN and ST14 is nafamostat, an anticoagulant drug which was shown to block SARS-CoV-2 infection in Calu-3 cells with an EC50 of 0.01 μM (54), Caco2 cells with an EC50 of 0.04 μM and VeroE6 cells with an EC50 of 23 μM (**Table 3**). This drug is also advanced to clinical trials in the RACONA study (NCT04352400) to test its efficacy in lowering lung function deterioration and reducing intensive care admissions in COVID-19 patients. Intravenous administration of nafamostat mesylate in a randomised controlled trial did not show evidence as an effective treatment for COVID-19 in a limited cohort of 42 patients; however, it was suggested for further investigation as an early treatment option for COVID-19 (55).

**Table 3.**
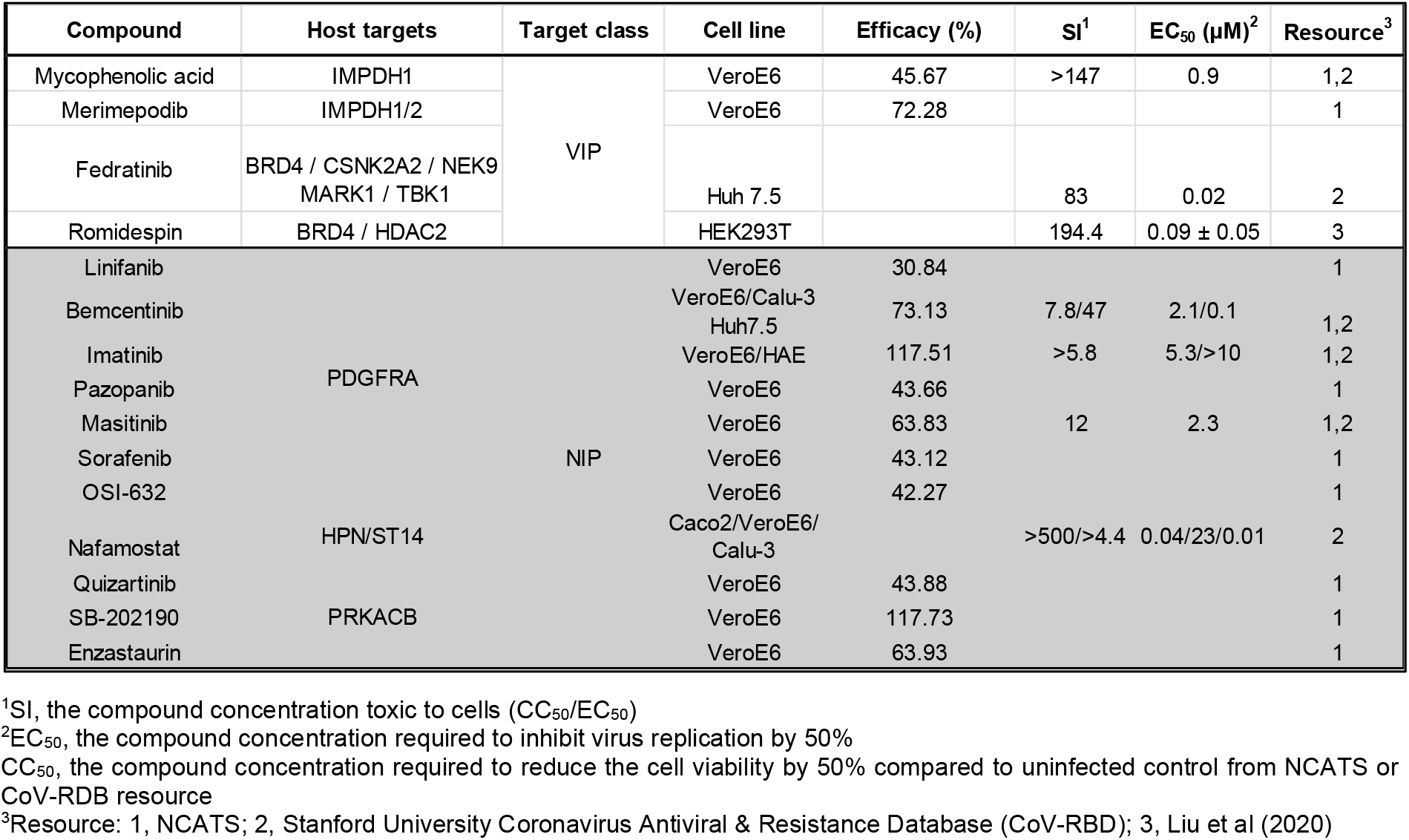
Compounds identified by our network-based approach and validated as antivirals in other studies. Supplementary Table S4 lists all the compounds identified in NCATS resource.

The IMPDH inhibitors mycophenolic acid (MPA) and merimepodib have previously been shown to inhibit SARS-CoV-2 infection in VeroE6 cells (56-58). MPA also showed antiviral effect in lung organoids and in nasal and bronchial epithelial explants (59,60). Specifically, the compounds are reported to have antiviral efficacies of 45.67% and a SI of >147 for MPA and efficacy of 72.28% for merimepodib in VeroE6 cells (**Table 3**). In our CPE assay, the compounds showed a modest degree of antiviral effect, and no apparent toxicity in 293TAT cells. More specifically, MPA showed a maximum antiviral efficacy of 39% at 0.01 μM (**Figure 7A**), and merimepodib showed peak antiviral effect of 26% at 0.01-0.03 μM (**Figure 7B**). However, there was no dose dependency in the observed antiviral efficacy. These results further demonstrate the cell context-specificity of the treatment effects, both proviral and antiviral, as measured by various experimental assays in different cell lines.

**Figure 7.**
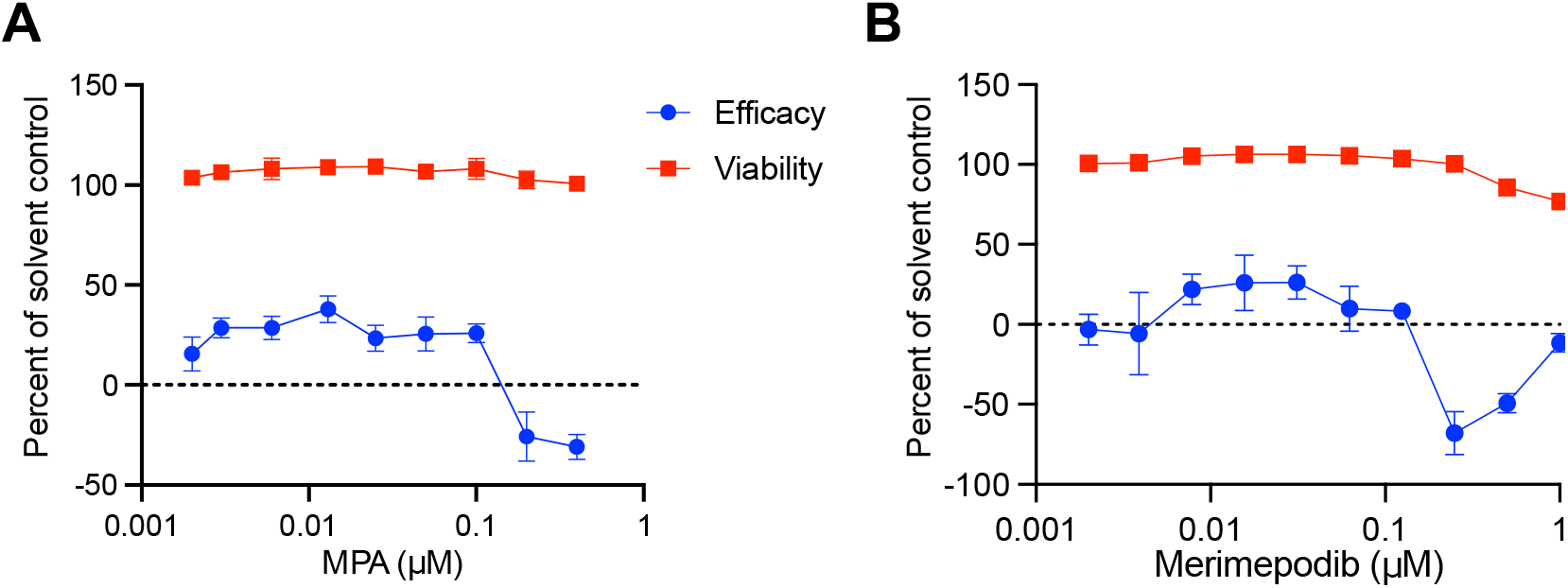
IMPDH inhibitors mycophenolic acid (MPA) and merimepodib show modest antiviral effects on SARS-CoV-2 infection in 293TAT cells. 293TAT cells were treated with the indicated concentrations of compounds immediately prior to infection with SARS-CoV-2 at a MOI of 0.01. Parallel wells contained cells treated only with compounds to study toxicity. Forty-eight hours post infection, cell viability was assessed using CellTiter-Glo and antiviral efficacy and viability were calculated as described in Materials and Methods. Data points reflect average and standard deviations of triplicate experiments per condition.

## DISCUSSION

Over the past two years, hundreds of predictive tools for COVID-19 diagnosis and prognosis have been developed based on the accumulated proteomic, transcriptomic, and clinical data sets. However, based on systematic literature reviews, most of the predictive approaches developed to support medical decision making suffer from training data of low quality and poor reporting of the models (61). For instance, so far none of the models for prediction of clinical outcome developed using chest radiographs and CT scans are of potential clinical use, mainly due to methodological flaws and high or unclear risk of bias in the imaging datasets used for the model training (62). Compared to supervised prediction models, network-based models provide a holistic view of the biological system being studied (here, virus-host interactions), and do not require large amounts of outcome data for model training. Network approaches can also broaden our understanding of the mechanisms of viral infections.

Consistent with previous studies, we showed that virus-interacting proteins (VIP) tend to be in essential positions for PPI network information flow. Gysi et al demonstrated that most of the drugs that successfully reduced viral infection do not bind the proteins targeted by SARS-CoV-2, indicating that these drugs rely on network- or pathway-level mechanisms that cannot be identified using docking-based strategies (13). We therefore expanded the target space to include additional host proteins that can be useful for disease pathogenesis, but may not be directly targeted by viral proteins. Along with cell context-specificity, it is important to consider interactions of the host proteins, as they can play pivotal roles in virus modulation. To elucidate the importance of the host proteins identified by the network approach, we screened for their effects on SARS-CoV-2 infection using selective inhibitors of the cellular proteins.

Our study identified antiviral proteins by virtue of the compounds inhibiting their targets acting as pro-viral agents. In particular, inhibition of epigenetic regulators BRD4 and HDAC2 by JQ1, vorinostat and romidepsin resulted in an increase in viral RNA, suggesting that these proteins function in an antiviral fashion (**Figure 6**). The viral protein Nsp 5, which targets HDAC2 (**Table 1**), cleaves the SARS-CoV-2 polyprotein and has developed strategies to inhibit the transport of HDAC into the nucleus and potentially affect the ability of HDAC2 to mediate inflammation and interferon response (14). Also, HDAC2 has been highlighted as one of the potential determinants of age-related susceptibility to SARS-CoV-2 infection, since it is downregulated in lungs of older individuals (63). HDAC2 and BRD4 interact with several other epigenetic modifiers in the SARS-CoV-2 network (**Figure 8**). HDAC2 erases acetyl marks on histones H3 and H4 to promote predominantly repression of transcription, and BRD4 is a histone acetyl reader that is predominantly an activator of transcription. Both proteins regulate a multitude of genes, including the antiviral response genes, and their inhibition is generally growth-suppressive, which may explain the pro-viral effect of their inhibition in the context of the SARS-CoV2 infection. Epigenetic modifications are significant in regulating cellular function, both in health and in disease. Moreover, BRD4 and HDACs have been shown to play diverse roles in virus infections, and viruses can evade various pathways involving these proteins for their gain (64,65). Chen et.al. showed that the viral protein E in its acetylated form can directly bind to the second bromodomain of BRD4, and it has evolved to antagonise interferon responses via inhibition of BET proteins (66).

**Figure 8.**
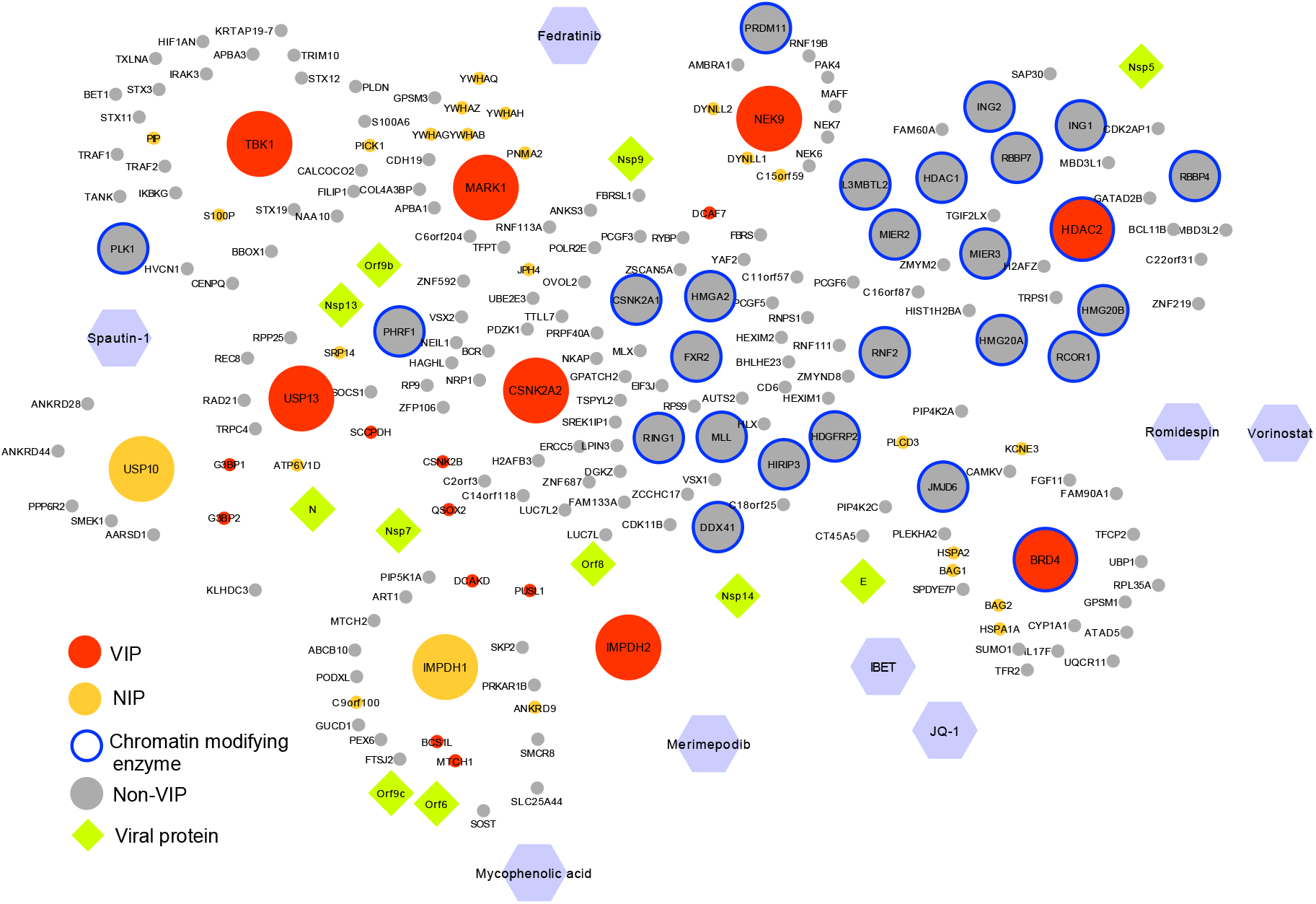
Network representation of selected targets and their protein-protein and compound-protein interactions with the validated compounds. Targets and their nearest neighbours are shown in the network.

Recently, Daniloski et.al. showed a pro-viral effect of vorinostat in human alveolar basal epithelial carcinoma cells expressing ACE2 (67). On the other hand, computational analysis using network clustering along with connectivity map (C-MAP) analysis of SARS-COV-2 PPI network identified vorinostat as potentially antiviral (68). Such contradicting results may be due to dependency of the compound’s effect on the cell context. To explore cell context specificity, we tested JQ1 and IBET151, two inhibitors of BRD4, in Calu-3 cells and observed that the compounds showed antiviral effects but with more pronounced toxicity in non-infected cells (**Figure S3**). Since the host and virus engage in an arms race in which the infected cell tries to inhibit the virus, and the virus tries to manipulate the host cell response for its own needs, modulation of BRD4 and HDACs should be further investigated for their ability to inhibit or enhance SARS-CoV-2. To further elucidate the underlying mechanism of HDAC, one could test the effects of HDAC and BRD4 agonists on SARS-CoV-2 infections.

The deubiquitinase proteins USP10 and USP13 may act as antiviral cellular proteins, since spautin-1, an autophagy inhibitor that targets these ubiquitinases, enhances SARS-CoV-2 infection. USP10 is a NIP that is involved in varied cellular processes, including DNA damage response, stress granule formation, and maintenance of intracellular protein homeostasis by recycling of cellular proteins (69). To combat viral infection, the host cell triggers the core autophagic machinery to launch an antiviral response. Several viruses, including herpes simplex virus, human cytomegalovirus and Kaposi’s sarcoma associated herpesvirus, can subvert the autophagic response indicating that viruses can manipulate autophagy for immune evasion (70). Furthermore, in-vivo and in-vitro studies have shown that SARS-CoV-2 inhibits autophagy in infected cells, suggesting the use of compounds that induce autophagy as an antiviral treatment strategy (71). The study by Shang et.al. showed that autophagy inhibiting agents were beneficial in reducing infection in the hACE2 transgenic mouse model (72). Furthermore, the expression of USP10 in human airway epithelial cells plays a vital role in regulating cystic fibrosis transmembrane conductance regulator (CFTR), a protein that helps mucociliary clearance and elimination of pathogens from the lungs, which is beneficial to patients with pneumonia (69,73). The roles of USP10 and USP13 in SARS-CoV-2 infection therefore warrants further investigation. Another protein that might be worth pursuing for some variants of concern (VOCs), such as Omicron that rely more heavily on the endosomal entry pathway (43), is RAB7A which was initially deprioritized for the experimental validations.

The potential drawback of direct-acting antiviral agents used as a monotherapy is the potential emergence of resistance. Inhibiting a host target alone is unlikely to have the potency one can achieve by inhibiting a viral protein such as a viral RNA polymerase. However, host-targeted antiviral drugs exploit the dependence of the virus on specific host proteins and pathways during replication, and hence drug resistance is less likely with the host-targeting antivirals. Combination therapies offer widespread well-documented advantages in the treatment of complex diseases such as cancer, HIV, and HCV (74-76). Since we observed weak antiviral activity for IMPDH inhibitors MPA and merimepodib, we further explored whether their activity could be improved by combination with molnupiravir (MPV), a directly acting antiviral (DAA) that targets the viral RNA dependent RNA polymerase (RDRP), and that has been approved for emergency use by the U.S Food and drug Administration (77,78). We evaluated the two-drug combinations against SARS-CoV-2 infections in 293TAT cells. The combinations had minimal synergistic effect, but importantly, there was no detectable toxicity in non-infected cells in either of the combination assays (**Figure S4**). Several other studies have evaluated the effects of combination therapy in SARS-CoV-2. For instance, nafamostat combined with MPV showed a significant increase in antiviral activity compared to their solo treatments in human nasal epithelial cells (79). Another effective combination reported in Calu-3 lung epithelial cells is between erlotinib and sunitinib, a drug identified by our network analysis (80).

In summary, we have demonstrated that a network-based approach that identifies SARS-CoV-2 VIPs and NIPs can be leveraged to prioritise host modulators of viral infection. Both compound sensitivity assays and protein-protein interactions are very context specific, hence it is important to consider their molecular characteristics in various cell lines to provide more consistent conclusions. Furthermore, testing of the compounds in non-infected cells is critical for avoiding broadly toxic compound responses, and to identify either safer small-molecules for therapeutic applications, or selective chemical tools to probe virus-host interactions that regulate virus infection. The model predictions could be further improved by leveraging compound– target binding interactions already in the network modelling, along with the gene expression datasets across different cell lines perturbed by the explored drugs, using the C-MAP approach.

## MATERIALS AND METHODS

### SARS-CoV-2 protein-protein interaction network construction

The SARS-CoV-2 protein-protein interactions (PPIs) were extracted from Gordon *et al*. (14). The PPIs include physical interaction among 26 viral proteins and 332 human proteins identified using affinity purification mass spectrometry in HEK293T cell line. We constructed the SARS-CoV-2 network by overlaying the HEK293T PPI network with the interactions from the BioPlex Interactome (15). The BioPlex PPI network includes 13957 proteins and ∼120,000 pairwise interactions. Out of the 332 viral-targeted host proteins, called viral interacting proteins (VIPs), 298 had an interaction in the BioPlex database.

For each of the 298 VIPs, we considered the protein interaction partners that were directly interacting with the viral-targeted proteins, that is, their nearest neighbours, thus creating an induced subgraph from the BioPlex HEK293T network. The final SARS-CoV-2 interaction network included 3978 proteins and 41015 interactions, which were used for further downstream analysis. This PPI network is context-specific, as we incorporated PPIs from HEK293T cell line that was used by Gordan *et al*. (14), along with additional proteins that are closest to the VIPs in terms of their connectivity/interaction to VIP proteins.

All the network visualisations were made in Cytoscape software (81), and the network topological parameters, such as degree connectivity and betweenness centrality, were computed using the igraph R package (82). The degree of a node is the number of interactions it has to the other nodes in the network, and betweenness of a node is a measure of centrality that corresponds to the number of shortest paths between each pair of nodes that pass through that node.

### Identification of host targets using protein prioritisation

We used a network-based protein prioritisation approach, the random walk with restart (RWR) (19), to identify host protein targets closest to the VIP protein target set. A random walk is a stochastic process, where the steps on the network take place with a certain probability. The process visits a sequence of adjacent nodes, forming a path, where at each step the next node is chosen at uniformly random among the neighbours of the current node. A RWR implements an additional restart probability, meaning that for every step taken in any direction, there is a positive probability of going back to the initial node. The advantage of the RWR method is that it provides a good representation of the topological similarity between the seed nodes and other nodes in the network. In the RWR method, each protein node in the network is ranked in the descending order based on the probability of them being visited by a random walker that starts from any of the seed nodes (here, the VIP nodes). The RWR process can be represented mathematically as:

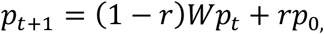

where *W* is column-normalised adjacency matrix of the network, *r* is the probability of restart, and *p*_*t*_ is the probability vector of being at node *i* at time step *t* to be visited by a random walker. *p*_*0*_ is the initial probability vector where the seed VIP nodes have equal probabilities and their sum equals to 1. We used Arete Cytoscape plugin that provides an exact solution, rather than an approximate or iterative solution for RWR, which is more robust to the restart probability parameter (*r* was set here equal to 0.7 which is the recommended default) (19,83).

### Biological characterisation of the network proteins

The GO and KEGG pathway enrichment analysis for the VIP and NIP sets was carried out using the clusterProfiler R package (84). The protein names were mapped to the NCBI entrez ID using the genome-wide annotation for human (org.Hs.eg.db) R package (85). The tissue expression of the proteins in the network were analysed using HPAnalyze (86), an R package that provides functions for retrieving, exploratory analysing and visualising the Human Protein Atlas data (27,28).

### Identification of potent compounds for the host targets

The predefined host target protein sets (VIPs and predicted NIPs) were queried in the ChEMBL database (26), a public bioactivity data repository, to find compounds that inhibit the identified host targets. A compound is active against a protein target when it binds to it and inhibits a specific molecular pathway. The strength of this inhibition can be quantified using several affinity measurements, such as IC_50_ and EC_50_, that refer to the *in vitro* concentration of the compound required to inhibit half of the target volume or produce half of the overall effect, respectively. The binding affinity that a compound has against a protein target can be defined by K_i_ (the inhibition constant) or K_d_ (the dissociation constant). These multi-dose bioactivity data (IC_50_, EC_50_, K_d_ and K_i_) were downloaded from the ChEMBL version 29 in PostgreSQL format and further processed in pgAdmin 4. We considered compounds with a bioactivity of less than 1000nM against the VIP or NIP targets as potent compounds for these host targets. In cases where multiple compounds inhibit a particular host protein, we selected the compounds with the highest potency (lowest bioactivity value) as inhibitors for experimental validations.

When comparing with the NCATS compound database, we converted the ChEMBL IDs into PubChem SIDs. The antiviral compounds were classified based on the curve rank, which is a numeric score, developed internally by NCATS, that combines classes of the dose-response curves and efficacy, such that more potent and effective compounds with higher quality curves are assigned a higher rank (87). In our analyses, all compounds with a curve rank >0 were classified as antivirals (88).

### Statistical analysis

All statistical analyses were performed using R (89) and Python (90). The two-sample Kolmogorov-Smirnov test was employed to compare the distributions of compounds and targets between the VIP and NIP sets, with the null hypothesis being that the distributions are similar, using the Scipy package (91).

### Cell lines, culture condition and compounds

Vero E6 and Calu-3 cells were maintained in standard medium (Minimum Essential Medium (MEM; catalogue number 11095; Gibco) supplemented with 9% foetal bovine serum (FBS; catalogue number SH3007103; HyClone) and 1% penicillin-streptomycin (catalogue number 15140; Gibco); 293TAT cells were maintained in DMEM (catalogue number 11995; Gibco), 9% FBS (catalogue number SH3007103; HyClone), 1% penicillin-streptomycin (catalogue number 15140; Gibco), 1% non-essential amino acids (NEAA; catalogue number 11140; Gibco), and 20mM HEPES (catalogue number 15630; Gibco). Compounds vorinostat (catalogue number S1047; Sellckchem), romidespin (catalogue number S3020; Sellckchem), spautin-1 (catalogue number S7888; Sellckchem), fedratinib (catalogue number S2736; Sellckchem), merimepodib (catalogue number S6689; Sellckchem), mycophenolic acid (catalogue number HYB0421; MedChem Express), (+)-JQ-1 (catalogue number HY13030; MedChem Express) and Molnupiravir (catalogue number HY125033 MedChem Express) were tested.

### Viral strain, infection and compound testing

Infectious SARS-CoV-2 was obtained from BEI Resources (Isolate USA-WA1/2020 NR-52281). 293TAT cells were plated the day prior to infection, while Calu3 cells were plated two days prior to infection. Compounds at various dose ranges were then added to plates and infectious SARS-CoV-2 was added to appropriate wells. Multiplicity of infection (MOI) of live SARS-CoV-2 is indicated in each figure legend. Mock-infected control wells received standard medium. Plates were then incubated at 37°C, 5% CO2 for 48 (293TAT) or 96 (Calu 3) hours. CellTiter-Glo (CTG) assay (Promega, G9243) was used to measure cell viability in virus infected and drug treated wells and in parallel mock-infected and drug treated wells. The assay measures the number of viable cells in culture by quantifying ATP, indicating the presence of metabolically active cells. Luminescence was read on a Biotek Synergy H4 plate reader. Each condition was conducted in triplicate.

Antiviral efficacy was calculated by comparing the relative light units (RLU) from virus infected (IFX) cells treated with drugs (IFXdrug) compared to infected cells treated with solvent control (IFXDMSO), and non-infected cells (nonIFX) treated with solvent control (nonIFXDMSO). Cell viability was calculated by comparing the RLU from non-infected cells (nonIFX) treated with drugs (nonIFXdrug) with non-infected cells treated with solvent control (nonIFXDMSO) using the following equations:

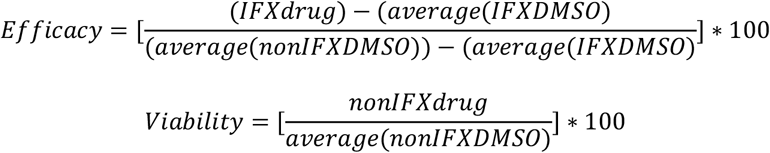

### SARS-CoV-2 qRT-PCR assay

RNA was isolated using the SingleShot Cell lysis kit (Bio-Rad). Briefly, at designated time points, cells were washed 1x with PBS followed by addition of a lysis buffer. After a 5-minute incubation at room temperature, liquid was transferred to a PCR plate and further incubated at 37C for 5 min followed by 5 min at 75C. The resulting RNA was used to synthesise complementary DNA (cDNA) using the iScript cDNA Synthesis kit (Bio-Rad). Briefly, 6 μL of RNA was mixed with 2 μL of reaction buffer, 1,5 μL of water, and 0.5 μL of reverse transcriptase. This was incubated for 30 min at 42C followed by 5 min at 85C. 10 μL of water was added to each product to dilute the cDNA. Quantitative RT-PCR was performed on a Roche Lightcycler 480 using SYBR Green (Bio-Rad) with the following primers (all primers listed in the 5’ to 3’ orientation).

human *RPS18*: TGC GAG TAC TCA ACA CCA ACA and CTT CGG CCC ACA CCC TTA AT

SARS-CoV-2 E: ACAGGTACGTTAATAGTTAATAGCGT and ATATTGCAGCAGTACGCACACA

For each reaction 2 μL of cDNA was mixed with 5 μL of SYBR, 2 μL of water and 0.5 μL of each primer. The RT-PCR reaction was 40 cycles of 5 seconds at 95C and 1 minute at 62C, followed by melt curve analysis. Melt curve analysis was used to assess whether single reaction products were produced. Expression was calculated relative to the housekeeping gene *RPS18*.

### Drug combination analysis

The two-drug checkerboard assays were carried out as before (92). The multi-dose combination experiment data were analysed in SynergyFinder v2.0 web-application (https://synergyfinder.fimm.fi), an open-source software for multi-drug combination analysis (93). We used the Bliss independence model to quantify the combination synergy. This model is based on a stochastic process in which drugs elicit their effects independently, and based on the probability of independent events, the expected combination effect can be calculated (94). The average Bliss synergy score over the entire dose-response matrix was calculated with SynergyFinder v2.0, along with the Maximum Synergistic Area (MSA), which is a 3×3 dose response area of the matrix, where the drug combination shows the highest synergistic effect. The selective efficacy was calculated by subtracting the toxicity (viability of mock-infected control cells) from the efficacy (inhibition of virus-infected cells). The selective efficacy quantifies the difference between efficacy and toxicity, meaning that a selective efficacy of 100 indicates that the drug combination inhibits the virus 100% and does not affect the mock-infected cells, while a selective efficacy of 0 indicates that the drug inhibits 100% of both the virus and mock-infected cells, indicating a high toxicity. The outliers for MPA at doses 3,6 and 12nM were corrected using DECREASE web-application (https://decrease.fimm.fi/) (95).

## Supporting Information

**Supplementary Figures:** Supplementary Figures S1-S4

**Supplementary Tables:** Supplementary Tables S1-4

## Acknowledgments

VR acknowledges the funding from the European Union’s Horizon 2020 Research and Innovation program under the Marie Skłodowska-Curie Actions Grant, agreement No. 80113 (Scientia fellowship). SJP was partially supported by the Department of Laboratory Medicine and Pathology (University of Washington). AF was supported by the Norwegian Research Council project number 237718 (BigInsight) and by NordForsk project 105572 (NordicMathCovid). TA was supported by the Academy of Finland and the Sigrid Jusélius Foundation. The following reagent was deposited by the Centres for Disease Control and Prevention and obtained through BEI Resources, NIAID, NIH: SARS-Related Coronavirus 2, Isolate USA-WA1/2020, NR-52281.

## Author contributions

VR, SP and TA conceptualized the study, supervised, and wrote the manuscript. VR performed mathematical modelling, analysed the data, and wrote the original draft. Jessica W performed the experimental analysis. PA performed the data analysis for the compound identification. AH performed the RT-PCR experiments. JS and SF designed the RT-PCR experiments and analysed the role of chromatin modifying enzymes. SP and JW conceptualised and designed the experimental analysis. AF conceptualised and supervised. AI performed the drug combination analysis. JS, SF, AH, AF, JW, TA wrote, reviewed, and edited the manuscript.

## Supplementary figures for

**Figure S1:**
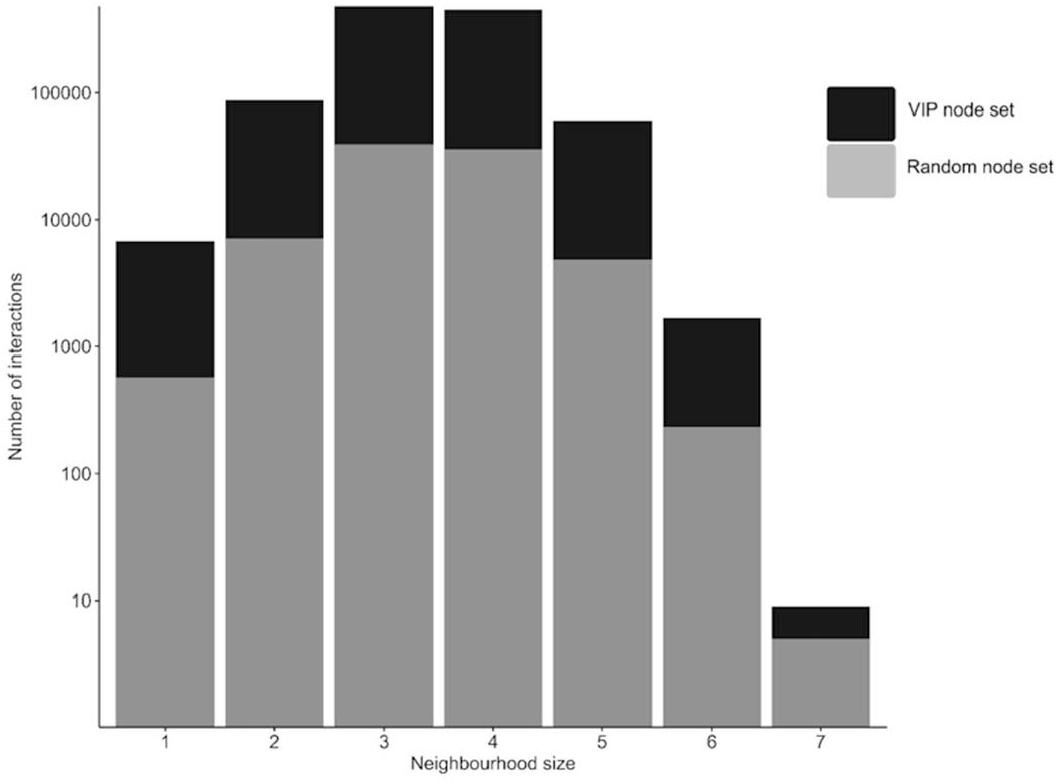
Neighborhood size (the shortest path distance) from VIP to non-VIP nodes compared to a random node set of the same size as the number of VIPs in the PPI network (297 VIPs).

**Figure S2.**
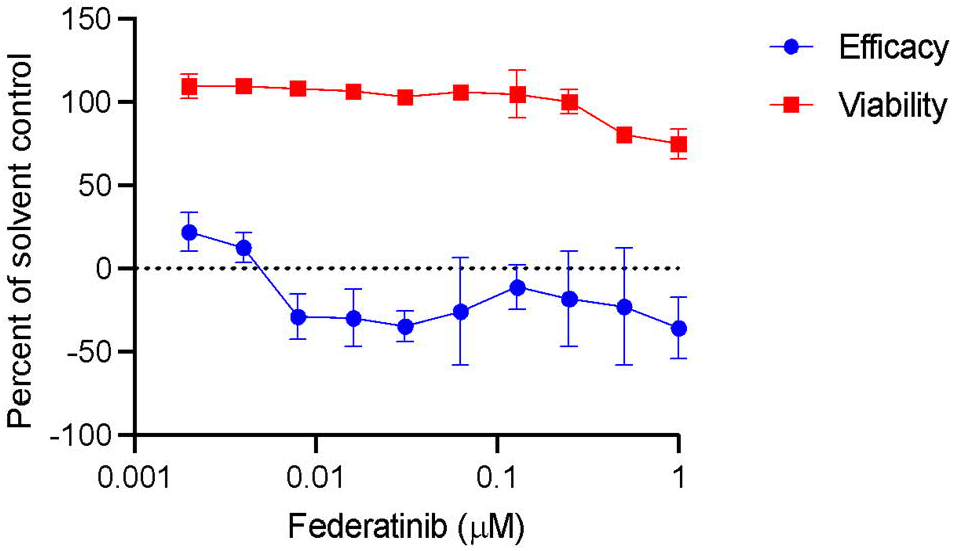
The effect of federatinib on SARS-CoV-2 infection. 293TAT cells were treated with the indicated concentrations of compounds immediately prior to infection with SARS-CoV-2 at a MOI of 0.01. Parallel wells contained cells treated only with compounds to study toxicity. Forty-eight hours post infection, cell viability was assessed using CellTiter-Glo and antiviral efficacy and viability were calculated as described in Materials and Methods. Data points reflect average and standard deviations of triplicate experiments per condition.

**Figure S3.**
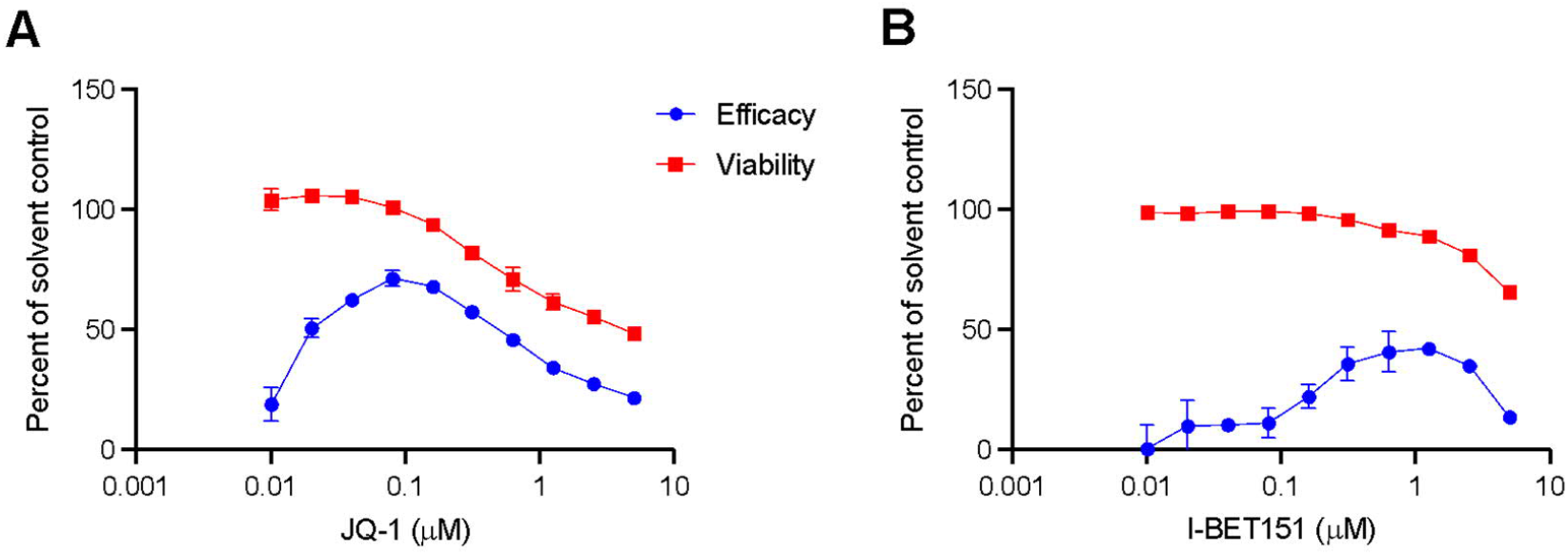
The effects of JQ1 and I-BET151 on SARS-CoV-2 infection. Calu-3 cells were treated with the indicated concentrations of compounds immediately prior to infection with SARS-CoV-2 at a MOI of 0.1. Parallel wells contained cells treated only with compounds to study toxicity. Ninety-six hours post infection, cell viability was assessed using CellTiter-Glo and antiviral efficacy and viability were calculated as described in the Materials and Methods. Data points reflect average and standard deviations of triplicate experiments per condition.

**Figure S4.**
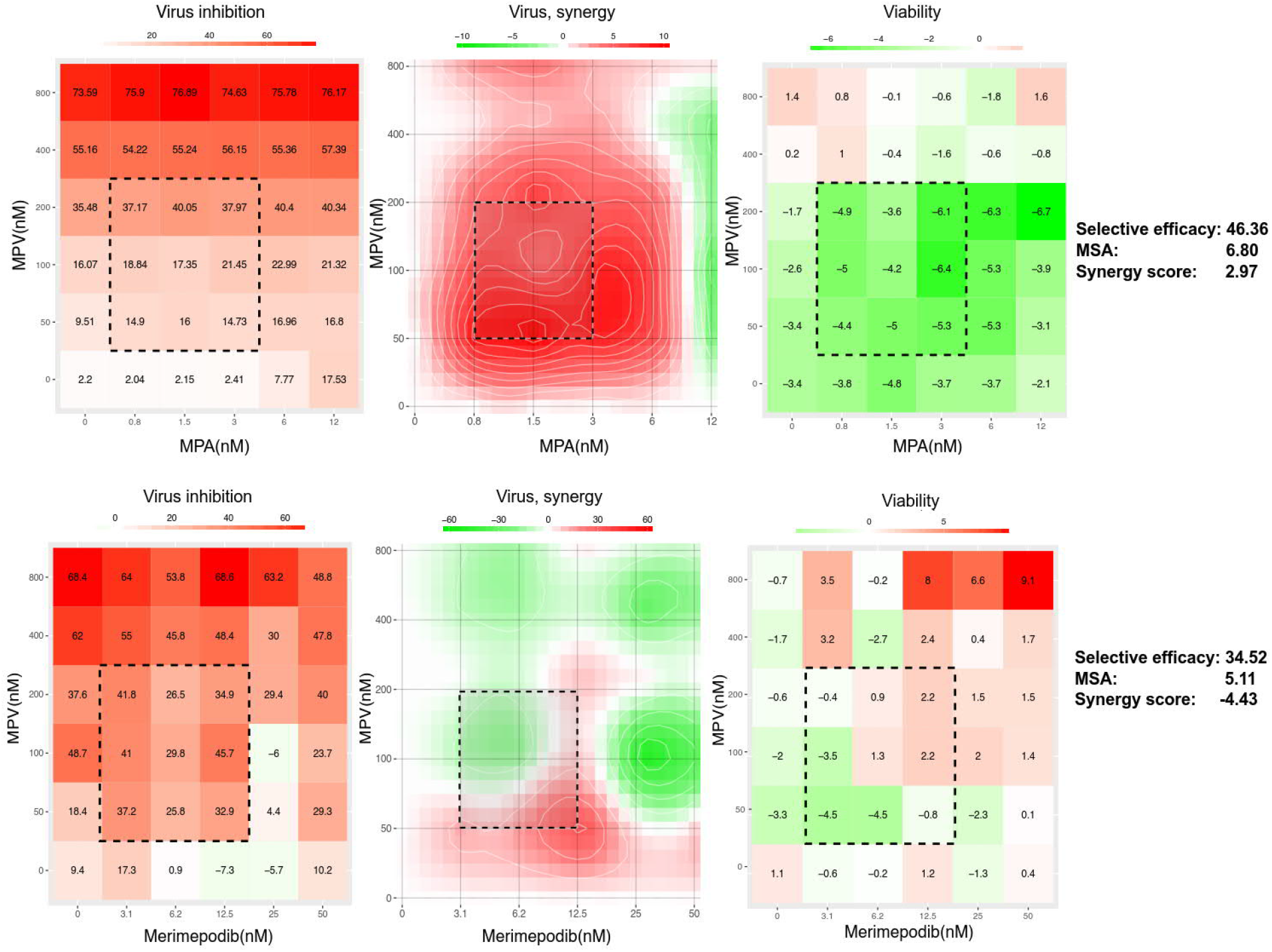
Dose-response matrix of SARS-CoV-2-infected 293TAT cells when combining molnupiravir (MPV, EIDD-1931) with either (A) MPA or (B) merimepodib. The maximum synergistic area (MSA, dotted-line square) and the Bliss synergy score were calculated with SynergyFinder v2.0, both for efficacy (virus infection) and viability (toxic effect on non-infected cells). Selective efficacy quantifies the difference in inhibition of virus-infected and mock-infected cells. A selective efficacy of 100 means that the drug combination inhibits 100% of the virus-infected cells and does not affect the mock-infected, drug-treated cells, while a selective efficacy of 0 means the drug kills both the virus and mock-infected cells. The selective efficacies of 46% and 35% for combinations of molnupiravir with MPA and merimepodib, respectively, indicate relatively high selective suppression of virus infection only. This is also seen in minimal co-inhibition of the viability of non-infected cells (right).

